# Thinking small: next-generation sensor networks close the size gap in vertebrate biologging

**DOI:** 10.1101/767749

**Authors:** Simon P. Ripperger, Gerald G. Carter, Rachel A. Page, Niklas Duda, Alexander Koelpin, Robert Weigel, Markus Hartmann, Thorsten Nowak, Jörn Thielecke, Michael Schadhauser, Jörg Robert, Sebastian Herbst, Klaus Meyer-Wegener, Peter Wägemann, Wolfgang Schröder-Preikschat, Björn Cassens, Rüdiger Kapitza, Falko Dressler, Frieder Mayer

**Affiliations:** Museum für Naturkunde, Invalidenstraße 43, 10115 Berlin, Germany; Smithsonian Tropical Research Institute, Apartado 0843-03092, Balboa, Ancón, Republic of Panama; Department of Evolution, Ecology, and Organismal Biology, The Ohio State University Columbus, OH, USA; Institute for Electronics Engineering, Friedrich-Alexander University Erlangen-Nürnberg (FAU), Wetterkreuz 15, 91058 Erlangen-Tennenlohe, Germany; Chair for Electronics and Sensor Systems, Brandenburg University of Technology, Siemens-Halske-Ring 14, 03046 Cottbus, Germany; Institute of Information Technology (Communication Electronics) LIKE, Friedrich-Alexander University Erlangen-Nürnberg (FAU), Am Weichselgarten 33 in 91058 Erlangen-Tennenlohe; Department of Computer Science, Friedrich-Alexander University Erlangen-Nürnberg (FAU), Martensstr. 3, 91058 Erlangen, Germany; Department of Computer Science, Friedrich-Alexander University Erlangen-Nürnberg (FAU), Martensstr.1, 91058 Erlangen; Carl-Friedrich-Gauß-Fakultät, Technische Universität Braunschweig, Mühlenpfordtstraße 23, 38106 Braunschweig, Germany; Heinz Nixdorf Institute and Dept. of Computer Science, Paderborn University, Warburger Str. 100, 33098 Paderborn, Germany; Berlin-Brandenburg Institute of Advanced Biodiversity Research, Altensteinstr. 34, 14195 Berlin, Germany

**Keywords:** wireless biologging network (WBN), ultra-low power, automated animal tracking, proximity sensing, bats, terrestrial biologging

## Abstract

Recent advances in animal tracking technology have ushered in a new era in biologging. However, the considerable size of many sophisticated biologging devices restricts their application to larger animals, while old-fashioned techniques often still represent the state-of-the-art for studying small vertebrates. In industrial applications, low-power wireless sensor networks fulfill requirements similar to those needed to monitor animal behavior at high resolution and at low tag weight. We developed a wireless biologging network (WBN), which enables simultaneous direct proximity sensing, high-resolution tracking, and long-range remote data download at tag weights of one to two grams. Deployments to study wild bats created social networks and flight trajectories of unprecedented quality. Our developments highlight the vast capabilities of WBNs and their potential to close an important gap in biologging: fully automated tracking and proximity sensing of small animals, even in closed habitats, at high spatial and temporal resolution.

## Introduction

Recent advances in animal tracking technology have ushered in a new era in biologging^1, 2^. By collecting data of unprecedented quantity and quality, automated methods have revolutionized numerous fields including animal ecology^3^, collective behavior^4^, migration^5^, and conservation biology^6^. For example, automated tracking of animals from space has advanced considerably over the past decade, in particular for observing large-scale movements^1^. However, satellite communication for localization or data access requires a lot of energy, and heavy transmitters greatly limit our ability to track smaller vertebrate species^1^. Efforts to further miniaturize increasingly powerful biologging devices culminated in the launch of the ICARUS initiative in 2019, which aims to achieve global animal observation at a small tag weight through a combination of GPS tracking, on-board sensing, energy harvesting, and energy-efficient data access from low space orbit^7^. ICARUS promises a great step forward in tracking large-scale movements such as migration. GPS tracking, however, is often not ideal or feasible for field biologists studying behavior on smaller spatial scales. GPS tracking of small vertebrate species is further limited by the considerable weight of GPS devices^1^. Satellite reception is hampered by complex habitats and impossible if animals go inside trees, caves, or underground burrows.

In industrial applications or for civilian surveillance, low-power wireless sensor networks (WSNs) fulfill requirements similar to those needed to track animal behavior at high resolution and at low tag weight^8^. Consistently, there have been numerous applications for WSNs in wildlife monitoring (‘biologging’) since the early 2000s^9^. In the last decade, more sophisticated approaches have created powerful monitoring systems, e.g., for high-resolution tracking^10^ and fully automated logging of social encounters^11, 12^. The major challenge in developing efficient wireless biologging networks (WBNs) is to design ultra-low power communication networks in order to maximize performance, minimize energy consumption, and reduce tag weight.

Here, we describe a system that takes WBNs to the next level: a multifunctional and thus <u>B</u>roadly <u>A</u>pplicable <u>T</u>racking <u>S</u>ystem (‘BATS’, Figure 1). We first present a solution for direct proximity sensing that enables the collection of proximity data at a temporal resolution of seconds, at tag weights of one to two grams, and with runtimes of up to several weeks depending on the sampling rate. Second, we describe an adaptive option for triangulating spatial positions based on Received Signal Strength (RSS) by ground-borne localization nodes. This adaptive option allows automated recording of robust movement trajectories even in structurally complex habitats. Third, we explore a new, almost energy neutral solution for remote data access over distances of several kilometers at low data rates. Finally, we present an energy model that shows the effect of the parameter settings of software tasks on the runtime of the animal-borne tag. First deployments of BATS have resulted in proximity and tracking data of unprecedented quality and have demonstrated the high potential of WBNs for studying (social) behavior. Our developments highlight the vast capabilities of WBNs and their potential to close an important gap in biologging: fully automated tracking and proximity sensing of small animals, even in closed habitats, at high spatial and temporal resolution.

**Figure 1:**
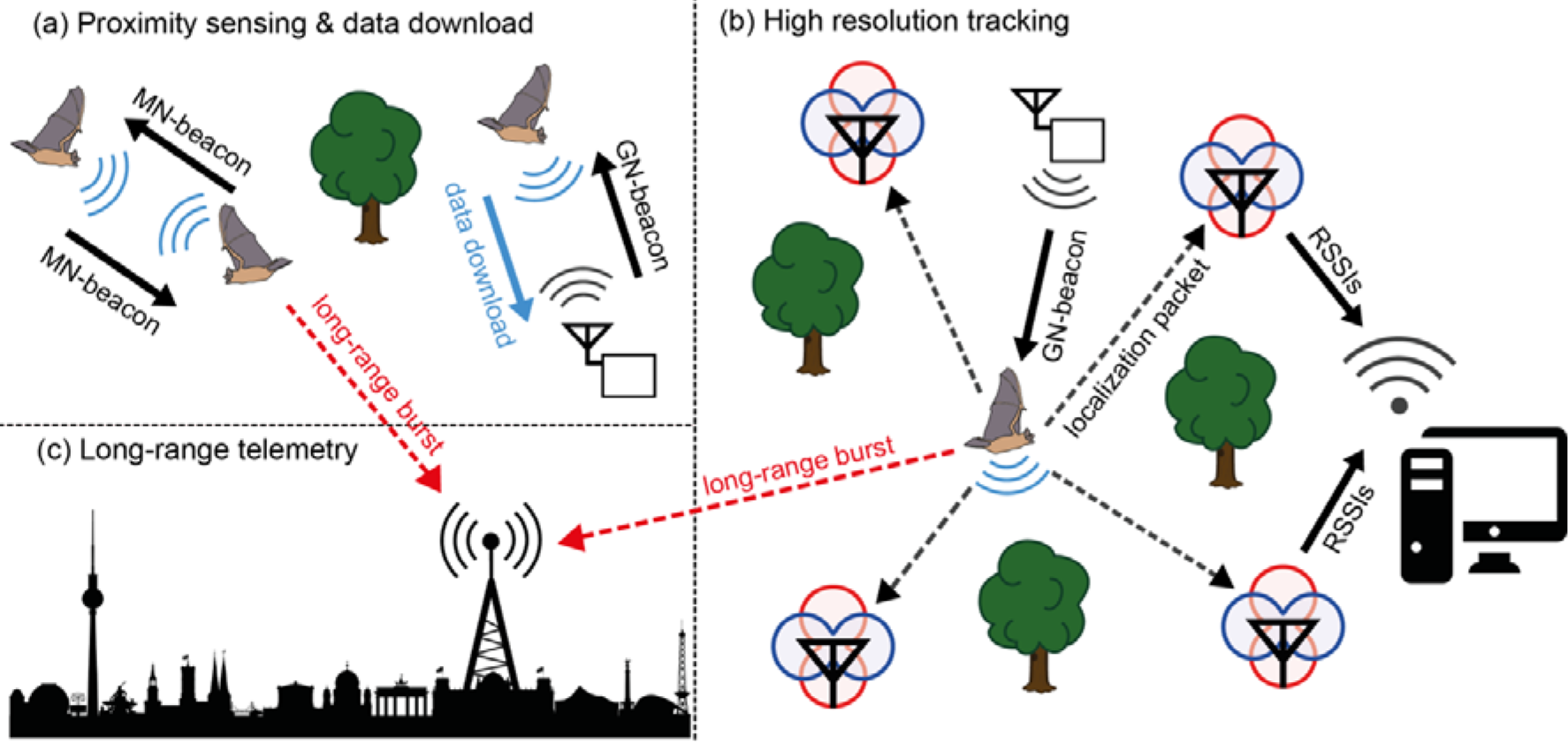
BATS overview. (a) Animal-borne mobile nodes (MNs) document animal-animal meetings, which are triggered by MN-beacons 24/7 and independently of ground infrastructure. Each mobile node forwards its meeting data when it receives beacons from a ground node (GN) that is dedicated to downloading and storing data. (b) When a tagged animal enters a grid of localization nodes (depicted by an antenna with red/blue gain patterns), a beacon of a tracking-dedicated GN triggers the transmission of localization packets from the MN to the localization nodes. Received signal strength indicators (RSSIs) of the impinging localization packets are then sent from the localization nodes to a work station via a WLAN. (c) Long-range bursts, which contain encoded sensor data, are received by long-range receivers. Long-range telemetry enables data transmission over distances of several kilometers at a low data rate.

## Results

The modular structure of BATS allows researchers to combine proximity sensing, long-range telemetry, and high-resolution tracking (Figure 1) depending on the research question and the behavior of the animals. We chose bats to test and validate the system since they are small-bodied and move fast in dense vegetation, both challenges to the performance of the WBN. Three recent field studies were conducted in temperate and tropical habitats on three bat species: greater mouse-eared bats (*Myotis myotis)*, common noctules (*Nyctalus noctula*) and common vampire bats (*Desmodus rotundus*). Each study documented high-resolution proximity data by direct proximity sensing among animals and automatically forwarding data to ground nodes (Figure 1a) that were deployed at roosting or foraging sites. Bat-borne mobile nodes that came within the reception range of the localization grid automatically increased their sampling rate to enable high-resolution localization (Figure 1b). Data for the synchronization of clocks was transmitted successfully over distances of more than 4 km by long-range telemetry (Figure 1c). The following sections describe the empirical validation of the system.

### High-resolution social network data from direct proximity sensing

Fifty individuals of one large natural colony of common vampire bats (*Desmodus rotundus*) were tagged simultaneously in Panama. Associations with other tagged bats are fluid and highly dynamic both during day and night. For example, Figure 2a shows the course of the meeting history and the dynamic range of degree centrality for a single bat (ID 56) over a two-day period. The high temporal resolution of meetings (all mobile nodes in reach communicate with each other every two seconds) also makes it possible to infer a behavior such as departure from the roost or movement within the roost. For example, foraging bouts can be identified by a sudden drop in meeting partners at night, which can be verified by contacts to ground nodes outside the roost. Autonomous direct proximity sensing allows monitoring changes in roosting associations, caused by moving among subgroups within the roost (Figure 2b, c) and it also allows inferring ‘social foraging networks’ outside the roost (Figure 2d). In addition, every meeting is labeled with a maximum signal strength intensity indicator (RSSI). This makes it possible to subset the meeting dataset according to signal strength, an estimate for proximity^13^. RSSI values can distinguish close-contact associations from associations based on merely occupying the same area.

**Figure 2:**
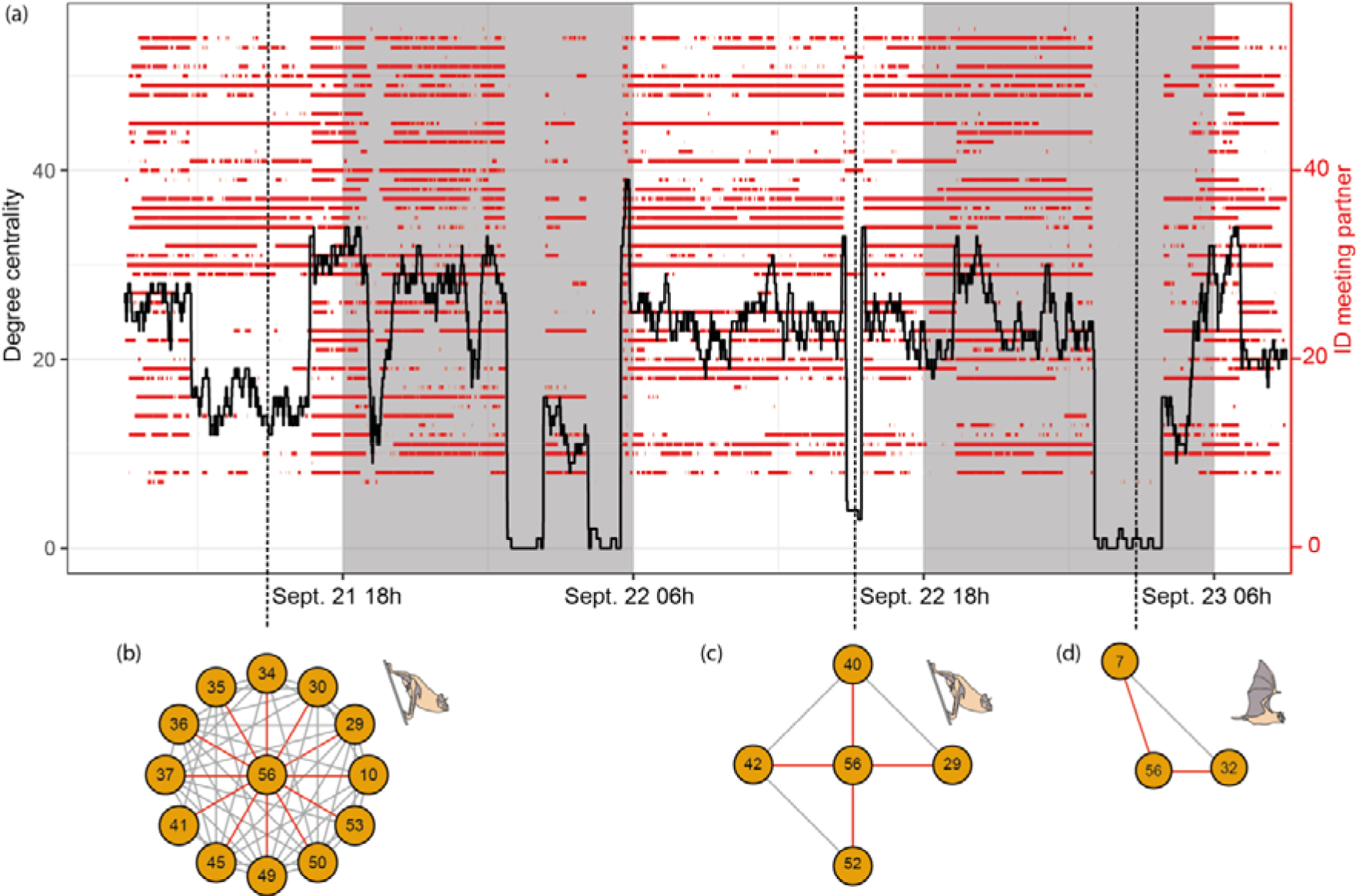
High-resolution association data in wild vampire bats. (a) Meeting history of a single vampire bat (ID 56; 50 tagged bats in total) with other tagged bats. Red lines show meetings between bat 56 and other tagged bats (right-hand y-axis). The black line shows the degree centrality (number of associated tagged bats, left-hand y-axis) of bat 56 every two seconds. Date and time are on the x-axis. Shaded areas indicate night time. Vertical dashed lines show egocentric social networks at each snapshot of time during roosting (b,c) and foraging (d). Associations with the focal bat are indicated by red lines.

The social networks created from direct proximity sensing are independent of the whereabouts of the tagged bats and provide an adaptive temporal resolution of seconds. Almost 400,000 individual meetings were recorded during the first eight days of our field test. To take bats as an example, the typical approach for collecting social network data has been to sample some unknown portion of co-roosting associations in a sample of identified roosts each day^14^. Our system allows complete networks of all bats every few seconds. This temporal resolution makes changes in social gathering directly visible if time slices in high-resolution data are small enough^15^. We believe this represents an extraordinary advance for studying such small free-ranging animals, and it allows for an analytical depth which is so far known predominantly from human social networks generated by communication among smart phones or social media^15^.

### RSSI-based localization from angle-of-arrival estimation

Seventeen wireless localization nodes were used to track tagged 11 mouse-eared bats (*Myotis myotis*) over an area of approximately 1.5 ha in an old, natural deciduous forest in northern Bavaria (Germany, Forchheim). We were able to reconstruct flight trajectories from foraging mouse-eared bats. Figure 3b shows as an example two trajectories of one foraging mouse-eared bat during two different nights in early August.

**Figure 3:**
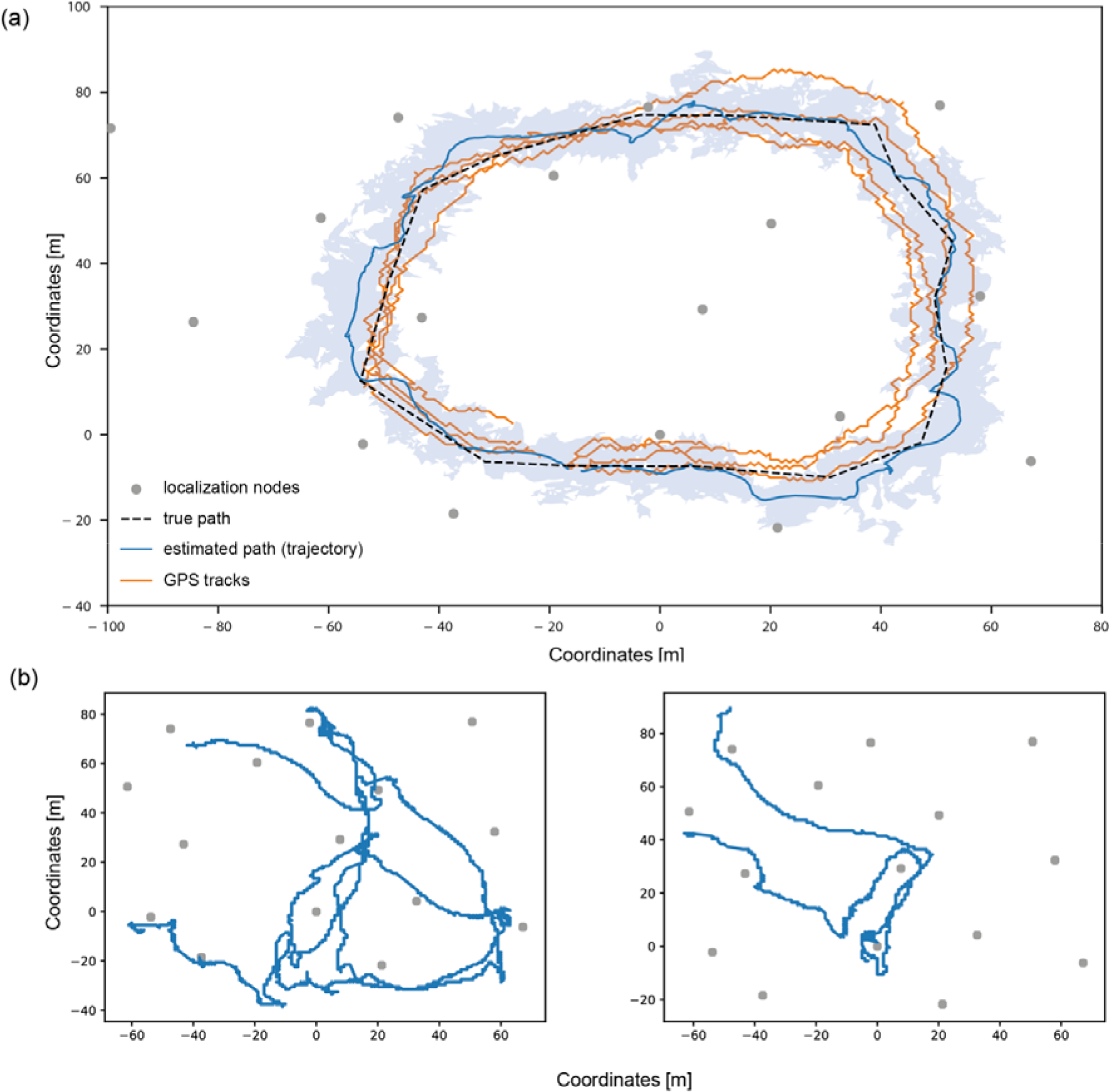
Tracking bat movements in a forest. (a) Tracking grid in a deciduous forest of Forchheim, Germany, consisting of 17 localization nodes (grey dots) covering an area of ca. 1.5 ha. Dashed black line: known reference path; blue line and blue shading: estimated path and average localization error obtained by BATS; yellow lines: four individual GPS tracks. (b) Estimated flight trajectories of a tagged mouse-eared bat during foraging on August 2^nd^ and 5^th^.

We evaluated the spatial resolution of the tracking system by estimating a trajectory from a defined reference path using unscented Kalman filters. The reference path and estimated trajectories are shown in Figure 3a. The trajectories were calculated from angle-of-arrival estimates of signals impinging on localization nodes. Angles were estimated from difference measurements of received signal strength (RSS) at two orthogonal antenna gain patterns. This procedure in combination with a set of post-processing techniques for probabilistic multipath mitigation makes the trajectories robust to multipath propagation. The calculated trajectory is based on 4,912 data sets, while one set was composed of up to 2×17 RSS difference measurements (one per frequency band), if all 17 localization nodes were within the reception range of the mobile node. For comparison, we also analyzed four tracks recorded by a 15 g heavy-duty Ornitela GPS tracker, which is commonly used for tracking large birds of up to 450 g body weight. The mean positioning error was 7.30 m for the Ornitela GPS tracker and 5.65 m for the trajectory of the BATS system.

We calculated the positioning accuracy at lower densities of the localization grid. Localization was less accurate with fewer localization nodes (Figure 4), but it was robust and comparable to the full tracking grid (17 nodes) when using 15 - 16 nodes. With 12 - 14 nodes, we observed increasing variation in average error rates. With 11 nodes, the mean error was similar to the results from GPS tracking. Variation increased steadily with lower numbers of nodes and the mean error reached more than 10 m with a maximum error rate of 34 m at 6 nodes. At such low grid densities, the localization results tended to diverge, resulting in increasing positioning errors. In addition, sparser grids lack robustness against multi-path scattering. Consequently, the node density may only be reduced to a certain point, while positioning errors remain quite stable (Figure 4).

**Figure 4:**
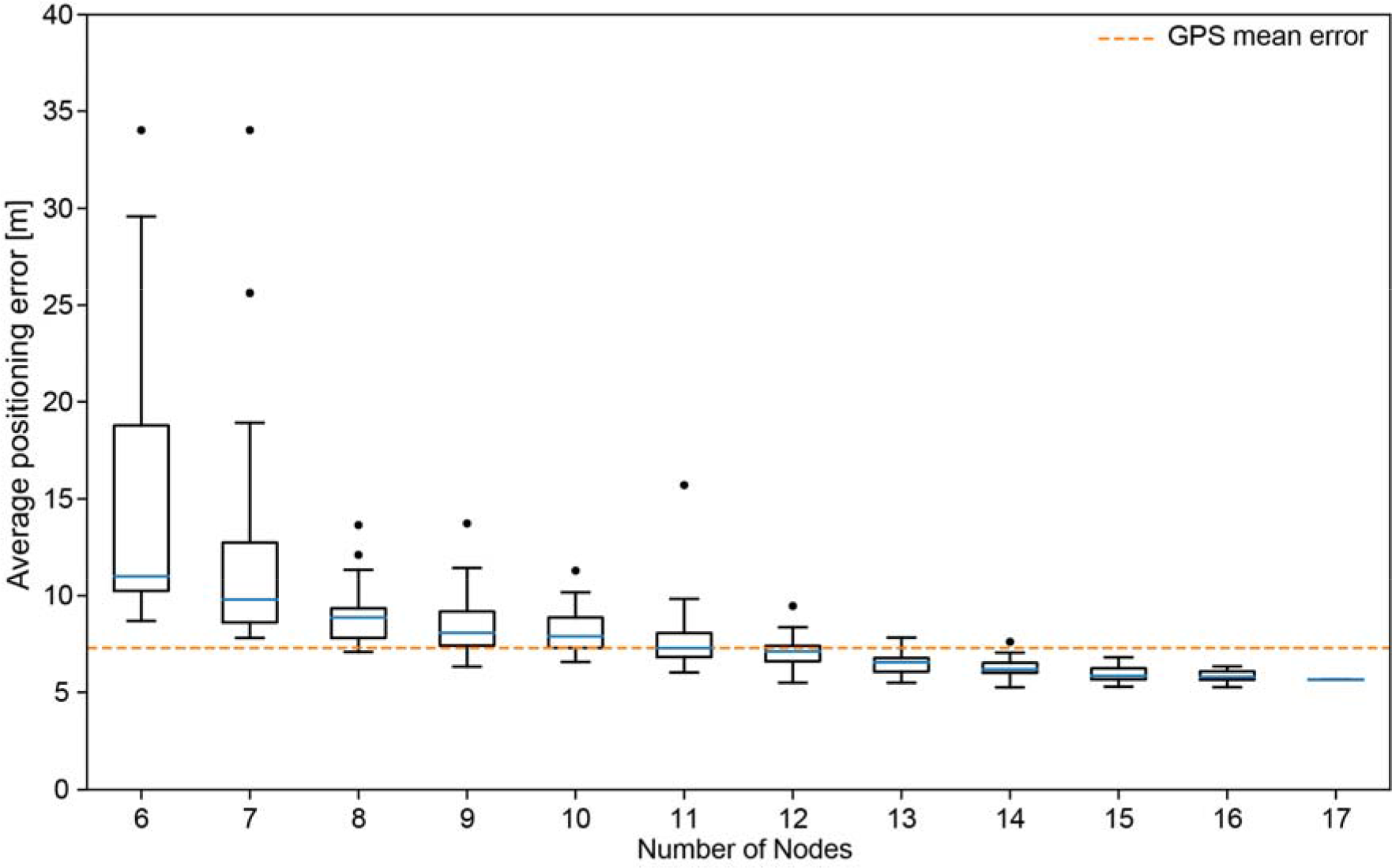
BATS tracking performance vs GPS tracking. Localization errors of a reference path of ca. 300 m by BATS are shown for different numbers of tracking nodes (6-17) in a deciduous forest of ca. 1.5 ha area. The average positioning error of four tracks of a heavy-duty commercial wildlife GPS tracker is shown for comparison by a yellow dashed line.

These analyses show that 11 localization nodes over an area of 1.5 ha in a forested habitat might be sufficient to construct high-resolution trajectories comparable in quality to a heavy-duty GPS tracker, which would only last for a few hours using a 15 g device, or to reverse GPS in open desert habitats^10^. Only moderate resources and human effort are needed to cover an area of a few hectares. For example, a setup as described above consisting of 11 localization nodes is deployed and configured by two people in one to two workdays.

### Distance verification of packet transmission by long-range telemetry

Transmission distances of data packages were measured in an urban green area. 34 noctule bats (*Nyctalus noctula*) were tagged in a forest within the city of Berlin and two long-range telemetry receivers were placed at a distance of about 1 and 4 km from the forest.

Mobile nodes transmitted with each MN-beacon a burst for long-range transmission of the mobile node’s time stamp. During a period of two weeks, we were able to receive more than 168,000 long-range bursts, which allowed us to successfully recover 9,511 complete timestamps from 32 individual bats. To mitigate the impairment by interfering transmission, one complete long-range telemetry packet is split over 24 single burst transmissions. At the long-range receiver, 24 subsequent bursts are merged to one actual long-range packet containing the ID and the bat’s time reference. This transmit scheme assures that the mobile node’s transmit module is only activated for a short time period avoiding stress on the batteries and hardware. In addition, it avoids interference of other channels by a time-frequency hopping pattern in transmission. Instead of a complete package loss, only a fraction of the collection of bursts might be corrupted, which can be reconstructed by means of error-correction codes at the receiver side. It is only due to this specialized telegram-splitting technique^16^ that a long-range transmission under extreme power-restrictions and vastly occupied frequency channels becomes possible.

Reception of long-range data should perform best when the tagged bats move in open airspace. However, we recovered a considerable number of these long-range bursts while bats were inside their roosts during the day. 563 long-range bursts received during the day were mapped to the known roosts of the bats, allowing us to measure the transmission distance. Fifty-seven long-range bursts from four bats inside their roost were recovered over distances of ca. 4.2 km (between roost 1 or 2 and the receiver at the cogeneration plant, Supplementary Figure 1). 506 long-range bursts from five tagged bats were recovered at distances between 667 and 819 m (between roost 1 or 3 and the receiver at the retirement home, Supplementary Figure 1). Burst retrieval over distances of more than 4 km was surprising. Theoretical calculations predicted transmission distances of about 5 km assuming barrier-free transmission^17^. In the field, however, signals had to pass first through the wooden wall of the tree roost and second the forest’s vegetation, which should greatly reduce transmission distance.

Data recovery is a major challenge in automated light-weight tracking-systems. Signal transmissions inherently suffer from limited transmission power under heavy losses due to distance, shadowing, and other interfering signals. Remote downlinks, e.g., per GSM (Global System for Mobile Communications), add considerable weight in the form of circuitry and battery^1^. Many light-weight trackers must therefore be retrieved, or energy harvesting must be used to counter-balance the expenses for remote data download^7, 18–20^, again adding weight for the required hardware components. When tagged animals move on predictable scales, energy-saving methods like transfer via VHF (Very High Frequency) or radio modems may be an option to receive data over distances of hundreds of meters to few kilometers^21^.

Our scenario explores options to decrease the energy expense for downloading stored data to a negligible proportion of the overall energy budget (compare Figure 5). For data download over short distances of ca. 100m, we accumulate and pre-process data on-board and use sophisticated communication protocols that maximize data-package reconstruction while minimizing energy demand^22, 23^. The above described long-range telemetry mode provides an option for robust transmission of small amounts of data at a low rate without the expenditure of additional energy due to the hybrid modulation of the signal. In comparison, other long-range systems for biologging, such as LoRa^12^, enable higher transmission rates. However, the bi-directional communication between transmitter and receiver strongly increases the energy demand no the mobile node.

**Figure 5:**
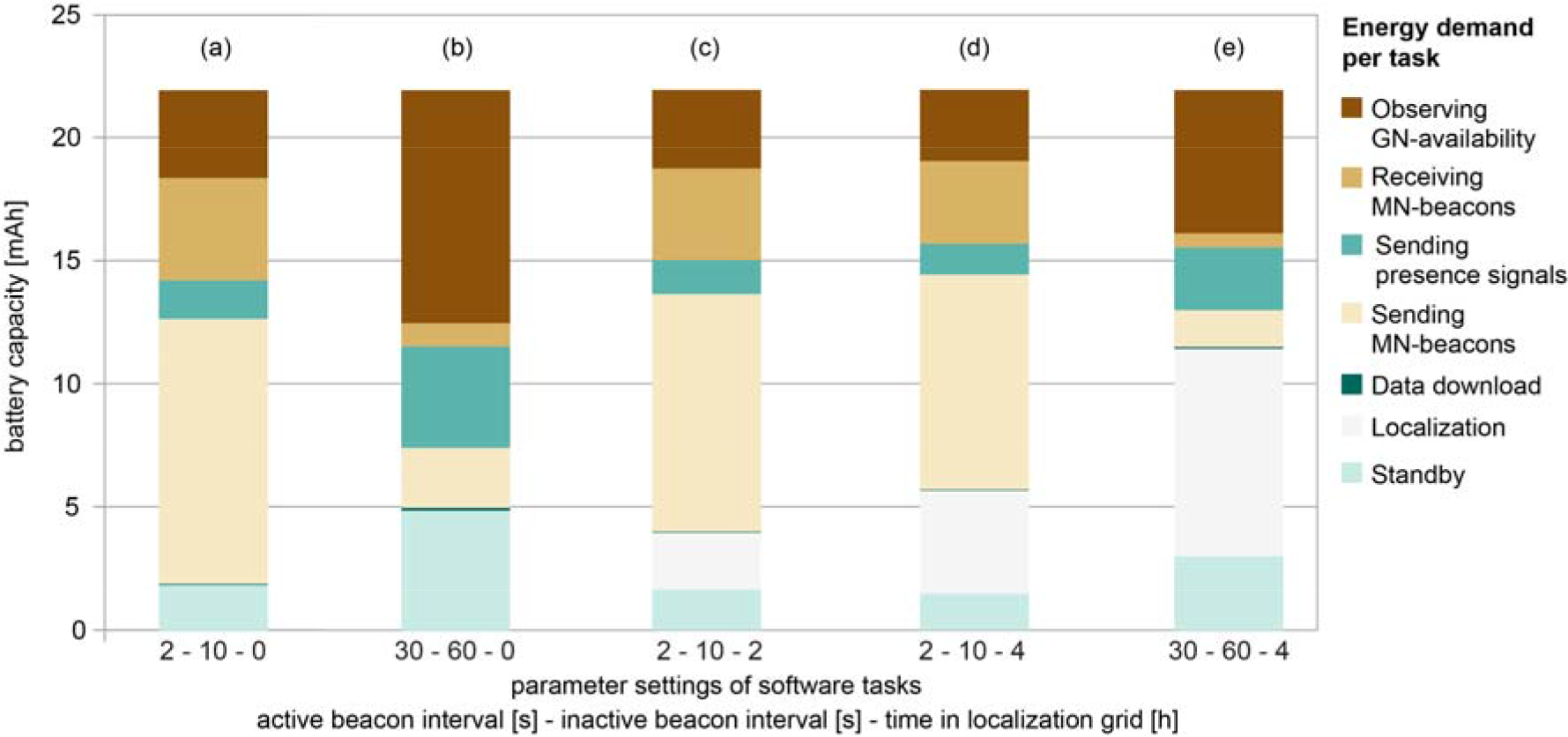
Energy distribution of software tasks of a mobile node powered by a 22mAh battery. Energy demand per software task depends on parameter settings for active/inactive beacon interval [s] and amount of time an animal spends in the localization grid [h]. The energy demand is shown for the seven major software tasks; MN = mobile node, GN = ground node. Zero time in the localization grid (a, b) refers to a pure proximity sensing scenario.

### Sensor node energy consumption and lifetime

A major strength of WBNs is the ease of adjusting parameters such as sampling rate and in turn energy consumption. These adjustments can maximize runtime for a given battery capacity, or alternatively maximize sampling rate to obtain higher resolution data. To investigate the impact of the different software-task parameters on runtime, we derived a model for energy consumption. We computed examples of runtimes of mobile nodes for two battery capacities and different parameter settings (Table 1). For example, the increase in energy consumption when tracking two to four hours per day can be compensated by extending the MN-beacon intervals. We achieve runtimes of at least five days using a 12mAh battery (corresponding to a 1g mobile node) even at the shortest beacon intervals of two seconds (active mode) and with two hours of high-resolution tracking per day. Depending on the parameter settings, we achieve runtimes of up to 13 days using the smaller battery and 25 days using 22mAh (Table 1).

**Table 1:**
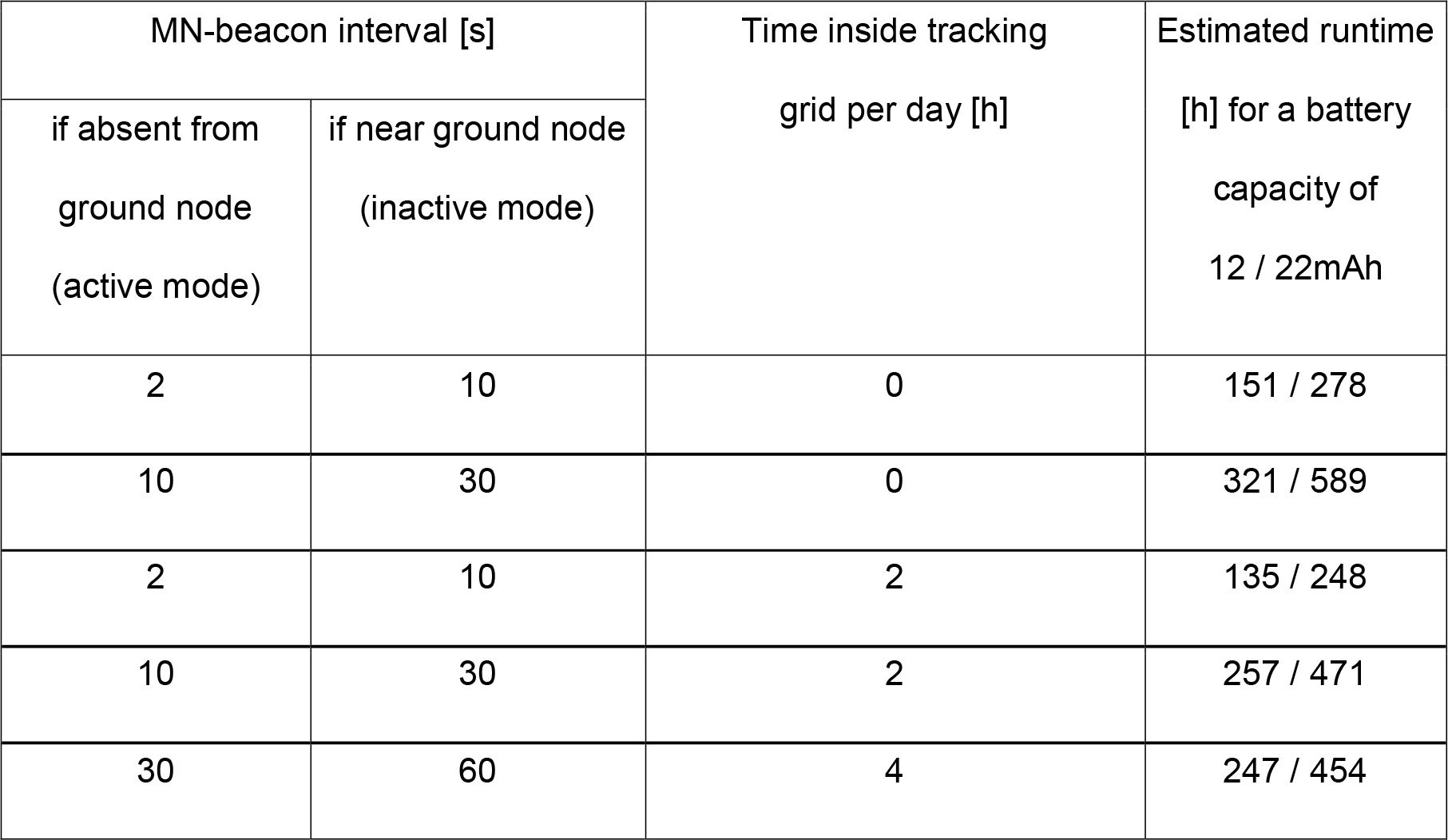
Estimated runtimes of mobile nodes for two battery capacities of 12 or 22 mAh inferred by an energy model for MN-runtime. While the model comprises seven energy consuming tasks, the shown runtimes are based only on varying MN-beacon intervals and localization time (i.e. animal is within the localization grid). For MN-beacon intervals two operation modes are possible, depending on whether an animal is within reception range of a ground node (inactive mode) or not (active mode).

| MN-beacon interval [s] |  | Time inside tracking<br>grid per day [h] | Estimated runtime<br>[h] for a battery<br>capacity of<br>12 / 22mAh |
| --- | --- | --- | --- |
| if absent from<br>ground node<br>(active mode) | if near ground node<br>(inactive mode) |  |  |
| 2 | 10 | 0 | 151 / 278 |
| 10 | 30 | 0 | 321 / 589 |
| 2 | 10 | 2 | 135 / 248 |
| 10 | 30 | 2 | 257 / 471 |
| 30 | 60 | 4 | 247 / 454 |

Figure 5 illustrates the energy consumption of the different software tasks on the mobile nodes of the six different scenarios described in Table 1. When localization is disabled (i.e., only proximity sensing), sending out beacons to wake up other mobile nodes to initiate meetings strongly drives the energy demand (Figure 5a, b). Therefore, modifying the MN-beacon intervals has the highest impact on runtime. When localization is enabled, tagged animals send localization packages whenever they enter the tracking grid. The high duty-cycle of sending localization packages (8/s) strongly decreases the runtime (Figure 5c-e, Table 1). At active/inactive MN-beacon intervals of 2/10s, a daily localization period of two hours decreases the overall runtime by 10.8% (Fig 5 a, c, Table 1). At four hours of localization, the energy demand for localization dominates the overall energy consumption, in particular at high MN-beacon intervals.

Model-derived runtimes were compared with empirical runtimes from field tests on noctule bats (*Nyctalus noctula*), where mobile nodes were either powered by a 12mAh battery or a 22mAh battery, resulting in MN weights of 1.1g to 1.9g depending on housing. The average runtime was 148h (max. 209h) for the small battery, while the model predicted 151h. For the larger battery, average runtime was 280h (max. 426h) with a predicted average value of 277h. For both batteries, the predicted values were very close to the observed (1.8% overestimation and 0.7% underestimation, respectively) indicating that the model is a reliable tool for experimental design of a field study.

## Discussion

In past years, developments in high-performance proximity sensing using significantly larger animal-borne tags^24^ and in ground-based high-resolution tracking in low-clutter desert environments^10^ have previously pushed the boundaries of what was technologically feasible. BATS takes the next step by combining these functionalities while keeping the tag weight at one to two gram. Adhering to the 5% rule^25^, even animals weighing as little as 20g can be tagged with this system. These smaller species make up a large proportion of birds and mammals (see Figure 3 in^1^) and WBNs will give researchers new capabilities to address a wide range of questions in animal behavior and ecology. Our adaptive and scalable system design provides great flexibility to tailor such a system, allowing adjustable use for any species- and study-specific requirement. The design of the mobile node allows to add multiple functionalities beyond the ones presented here, such as accelerometers, magnetometers, or even an on-board electrocardiogram (ECG) sensors^26^.

An early example of automated tracking of small-bodied animals on a limited geographic scale was ARTS, a system for automated VHF tracking, which was installed on Barro Colorado Island, Panama^27^. This six-year endeavor in the 2000s already highlighted the promising opportunities offered by WBNs: scalability, remote reconfiguration, full automation, and low-cost tags. Yet, a rather low positioning accuracy (~50 m) and restricted coverage were limiting factors^27^. Most of today’s solutions for automated tracking of small animals such as songbirds, bats, or rodents perform best at larger geographic scales or in open habitats. For example, current versions of 1 g GPS loggers are suitable to explore seasonal large-scale movements^18, 20^. However, they cannot reconstruct flight paths in a complex environment, since they can only collect around 100 fixes. Furthermore, the physical devices must be retrieved for data recovery, and satellite reception suffers within vegetated areas. Alternatively, reverse GPS can track small animals much more energy-efficiently and at much higher temporal and spatial resolution by measuring time-of-flight at ground-based receiving stations^10^. However, time-of-flight measurements are inherently affected by vegetation and perform best in open areas. Therefore, we combined signal strength measurements (including angle of arrival (AoA) estimates) from two frequency bands and probabilistic multipath mitigation^28^ to create a system that is robust to multipath propagation and thus performs well in complex environments. Common but costly measures to resolve multipath propagation are large-aperture antenna arrays for AoA tracking or large signal bandwidth for time-of-flight tracking. However, the future of animal tracking will most certainly center on low cost, ultra-low power integrated circuits, which are currently experiencing a noticeable push due to their broad applications in Industry 4.0 and 5G. This technology has the potential to dramatically boost the capabilities of biologging devices.

Contact networks of small-bodied animals have received increased attention in past years and are most commonly built from RFID- (radio-frequency identification) or PIT- (passive integrated transponder) tagged animals that were observed to be feeding or sleeping at the same site at the same time^29–31^. Later developments for direct encounter logging were able to log association independently of the locality. However, these sensors were either quite large and heavy^11^ or had short runtimes of less than 24h^32^ due to the high energy demand for the permanently active receiver — a major shortcoming for applications in small-bodied species. We show that the use of wake-up receivers and adaptive operation paired with novel wireless communication protocols dramatically reduce the energy demand of such wireless sensor tags. We believe that direct encounter logging or more precisely proximity sensing will enable diverse research in the future, as this approach creates large datasets, with additional sensor data providing the behavioral context closing the gap between social patterns and their underlying processes^2^.

Ongoing work on ultra-low power sensor networks not only targets animal tracking systems but a variety of “Internet of Things”-solutions in general. Energy efficiency is not only a question of hardware circuit design but also of how to interact across all relevant layers in the node’s software stack (i.e., application, and operating system). On the mobile node, the interaction aspect between communication layers (e.g., application and MAC layer) concerns the placement of certain functions (e.g., retransmissions) within the entire software hierarchy^33^ such that energy-efficient operation is not affected by unnecessary functional redundancies. Besides, cross-layer designs that optimize the timing of communication processes and make them deterministic at least within limits^34^ form the software-engineering basis for an overall energy-aware systems approach. In our scenario, energy-efficient and reliable communication between nodes are cross-cutting concerns, since failed communication attempts lead to additional overhead for retransmissions. Very promising examples for low-power communication initiation are novel selective wake-up receivers^35^, which allow the small tags to enter sleep modes in the nano-ampere range rather than constantly operating in the micro- or milli-ampere range. Selective wake-up concepts allow waking up dedicated recipients of a message (or a selected subset thereof) instead of waking up all systems in communication range. Integrated into the animal tracking nodes, this could enable the next quantum leap on low-power operation. The alternative to making the receiver operate on a lower energy budget is to make the communication more reliable. Recent advances in integrating coding for forward error correction into such lightweight systems show very promising results, e.g., using erasure codes^22^. Ultra-reliable communication protocols currently used in 5G networks can also be applied to localization nodes, which are used for quasi-live tracking in the bat tracking scenario. For example, the ground network can be used as a distributed antenna array, which allows the use of smart decoding algorithms for very weak communication signals to further optimize data recovery^36^.

## Conclusions

There is no single best method for tracking animal behavior. RFID- and PIT-tags allow monitoring presence of animals at known sites at low cost. Satellite-based localization will remain the method of choice to monitor large-scale movements such as migration or to explore unpredictable events such as nomadism^1, 37^. However, we believe that WBNs like BATS will greatly benefit biologging of small animal species that move over smaller and more predictable spatial scales, especially inside of habitats where signal transmission is constrained. The homologies of applications between mobile communications and biologging (e.g., Bluetooth low energy for communication among mobile nodes^12^) will boost the development of WBNs. Experimental setups including automated triggers (e.g., acoustic playbacks or other sensory cues) might easily be integrated and direct proximity sensing will bring exciting research opportunities. Such setups will allow to study the effect of social network dynamics on phenomena such as transmission of social information^30^ and pathogens^38^, and key ecosystem functions such as pollination and seed dispersal^39^.

## 4. Methods

### *The* WBN *hardware, software, and functionality*

Figure 1 is a schematic overview of BATS. In order to study its performance, we empirically evaluated the three major functions of the system: proximity sensing (Figure 1a), high-resolution tracking at local scales (Figure 1b), and long-range telemetry (Figure 1c). See Glossary (Supplementary Table 1) for definitions of terms.

### Proximity sensing

Any given mobile node (MN) dyad generates meetings whenever it comes within reception range (5 - 10 m depending on the environment). The animal-borne MN consists of a 22mm × 14mm Flex PCB circuit board, which is populated with a central System-on-Chip (EFR32, Silicon Labs) containing an ARM Cortex-M4 core and two radio frontends for 868/915MHz and 2.4GHz (Supplementary Figure 2). The transmitter in the sub-GHz frontend periodically sends MN-beacons, a signal that contains a wake-up sequence. The rate of beacons is configurable (see below). A low-power wake-up receiver on the MN triggers the conventional receiver to receive incoming information on the ID whenever a MN-beacon is received from another MN. Subsequently, a meeting is created between the communicating MN-dyad (Figure 1a left). While a conventional receiver draws a relatively high current in receiving mode waiting for incoming packages, a wake-up receiver achieves this functionality with a low current (yet, at cost of sensitivity and performance). When no further MN-beacons are received from the meeting partners for 5 MN-beacon intervals, the meeting is closed and stored to memory along with the ID of the meeting partner, meeting duration, maximum RSSI and a relative timestamp. The mobile node contains both persistent and volatile random-access memory for data storage.

The conventional receiver of the sub-GHz frontend is periodically activated to observe the presence of a ground node (at a fixed interval of every two seconds), which is indicated by a ground node beacon (GN-beacon), periodically broadcast by the transmitter of each ground node (Figure 1a right). The transmitter supports several configurations defining the main purpose of the ground node (GN) and enabling location-dependent adaptive operation of the WBN. (i) A download-dedicated GN broadcasts a signal that enables transmitting MN data based on a customizable RSSI threshold received at the mobile node. (ii) A tracking-dedicated ground node positioned within the grid of localization nodes for high-resolution tracking broadcasts a signal that activates the 2.4GHz frontend in addition to the sub-GHz front end on the MN, transmitting ‘localization packets’ at a rate of 8 packets per second. (iii) A presence-detection-dedicated GN triggers the transmission of ‘presence signals’ by an MN and stores incoming signals which can be used to determine presence/absence of tagged individuals (presence at resources or at sleeping sites). Combinations of functionalities (i-iii) may be used in a single GN if desired (e.g., a tracking-dedicated GN can also trigger data download). Incoming MN data is received by the GN and stored by a Raspberry Pi (Raspberry PI Foundation, Cambridge, UK) to a SD card along with the ID of the transmitting MN and the receiving GN, respectively, and a timestamp, which is provided by a GPS unit. The Raspberry Pi also hosts a WiFi allowing the user remote data access.

Visualization of proximity sensing data is facilitated by the custom-made software ‘meeting splitter’ (see Figure 2). For each specified mobile node ID, the current meeting partners are projected onto a discrete time axis (one second resolution). We specified a configurable time window around each point on the time axis (five seconds in case of Figure 2). All ongoing meetings, which overlap with the window around the respective point in time, are included in the set of associated bats at this particular point in time. The result per bat is a set of associated bats per each second in the dataset. A subsequent automated analysis classifies each meeting as inside or outside the roost, depending on the number of simultaneous meeting partners (not applied in this manuscript).

### RSSI-based high-resolution tracking

Localization nodes (LN) perform field strength measurements, which are collected by WLAN and are processed by a PC including a file system whenever animal-borne mobile nodes enter the localization grid (Figure 1b; localization nodes collect localization packets from mobile nodes; the transmission is triggered by a ground node). Each localization node comprises a software-defined radio (consisting of a radio-frequency frontend, a highly integrated analog-to-digital converter, a field-programmable gate array and a microcontroller) and two receiving antenna gain patterns each with two main lobes (Figure 1b, red and blue pattern, respectively). The bi-lobed shape indicates the directional sensitivity of the antenna, while the direction of each lobe represents its maximum in sensitivity. The red pattern is rotated by 90° compared to the blue pattern, and both are simultaneously used to estimate the angles of arrival (AoA) of the localization packages transmitted by the animal-borne MNs. The difference in received signal strength (RSS) of the two patterns relates to AoA: If the difference – RSS of the blue pattern minus RSS of the red pattern – is maximum, the wave front impinges on the localization node either from east or west; if the RSS difference is minimum, the direction of arrival is N or S. Accordingly, there are four options for the AoA if the difference is zero: NE, NW, SE or SW. These ambiguities are resolved by fusing measurements of several localization nodes.

This design allows us to exploit not only error-prone absolute field strength measurements^40^, but also fail-safe angles of arrival. These are not affected by faulty propagation laws or shadowing effects, because both error sources disappear when forming the RSS differences. The angular resolution of the AoA estimates improves with increasing ambiguity of the antenna pattern designs. However, more localization nodes have to be in reach to resolve the ambiguity^41^. During the Forchheim field trial, we collected up to 272 angle estimates per second when all 17 localization nodes were in reception range.

To further improve localization accuracy, we exploited three sources of information ((a) model-based Bayesian positioning, (b) frequency diversity, (c) retrodiction), which increase robustness against multipath propagation. This effect complicates the positioning process, in particular in structurally complex environments since wave fronts impinge a localization node out of the different directions of multiple reflectors (e.g., surrounding vegetation). The information sources to counteract multipath-related adverse effects are described in the following:

#### (a) Model-based Bayesian positioning

Due to the nature of multipath propagation, a stochastic model can be devised to characterize the resulting spread in the AoA estimates^28^. This AoA measurement model can be incorporated into the likelihood function of the recursive Bayesian positioning process, e.g., based on a Kalman filter or a related grid-based estimation filter^42^. The recursive estimation process yields a probability distribution characterizing where the bat may be, considering propagation characteristics from a local channel model^43^. All measurements are fused during the recursive process taking into account a movement model reflecting the flight characteristics of a bat (e.g., max. flight speed). The better the agreement of the various AoA estimates, the more pronounced the positioning probability distribution.

#### (b) Frequency diversity

Multipath propagation leads to frequency-dependent fading. We therefore measured field strength not only on the primary far-reaching carrier frequency at 868 MHz, but also on a secondary carrier frequency at 2.4 GHz. On both carrier frequencies, wave forms comprising several subcarriers are employed to enhance the field-strength based AoA estimation process. Due to the large carrier frequency separation (> 1.4GHz), frequency-dependent fading effects are de-correlated even if multipath time-of-flight differences are minor, i.e., in the range of a few meters, which corresponds to our accuracy level.

#### (c) Retrodiction

If we do not have to estimate the position of a particular bat in real-time, we can exploit all measurements of a bat to estimate a complete trajectory. Forward-backward filtering enhances estimation quality considerably, yielding a positioning quality in the range of 4 m (1 − σ). Performance limits of field-strength based positioning have been discussed in depth^41, 44^.

We evaluated the trade-off between tracking grid density and localization quality for the Forchheim setup, which comprised 17 localization nodes. In particular, we asked: how many localization nodes are required to obtain localization quality comparable to heavy-duty GPS tracking? We selected subsets of 6 to 16 out of the 17 localization nodes in order to observe the decrease in positioning accuracy with decreasing grid density. Trajectories including standard deviation were estimated for each subset of localization nodes. 17 configurations were calculated for the grid consisting of 16 nodes (all possible subsets of the full grid) and 25 unique, randomly chosen subsets for all remaining grid configurations (6 to 15 nodes, respectively) to obtain average errors for the given number of nodes (see Figure 4).

### Long-range telemetry

Our long-range telemetry approach aimed at transmitting long-range bursts from mobile nodes over distances of up to several kilometers – much longer distances than our download-dedicated ground nodes would allow for – within the city of Berlin, under harsh shadowing by obstacles (vegetation, buildings, etc.) or in presence of numerous interferes. We periodically transmitted ‘long-range bursts’, i.e., relative timestamps in form of seconds since mobile node start-up generated by a simple clock counter. These timestamps are crucial for post-processing of meeting because they allow accounting for clock drift on the mobile node. We embedded the long-range functionality into the existing modulation scheme using a hybrid phase-alternating modulation on top of the pure amplitude-modulated wake-up sequences of the MN-beacon^45^. As a consequence of the extreme energy limitation of the MN, we ensured the required Signal-to-Noise-Ratio (SNR) by counterbalancing the rate and the desired transmission distance. The combination of the hybrid modulation, the channel encoding procedure^17, 45^ and the ‘Telegram-splitting’ technique^16^ enables an ultra-low power long-range transmission without additional expenditure of energy. The long-range bursts were received at two long-range receivers, which were deployed on exposed sites (rooftops) at distances of ca. 200 - 1,800m (retirement home) and 3,300 - 4,500m (cogeneration plant; see Figure S1) to the proximate respectively ultimate border of the urban forest where the roosts of the tracked bats were located.

We quantified communication distances in the field, which was possible when a tagged bat occupied a known roost and was simultaneously received by the long-range receiver. We therefore matched timestamps of signals received simultaneously by GNs at roosting sites and at long-range receivers. In case of a match, we quantified the distances between roosts and long-range receivers in the R package geosphere using the Haversine function^46^. The empirically assessed communication distances have then been compared to a theoretical model of long-range transmission distances^47^. This model evaluates achievable rate and distance of transmission based on the energy relation of the SNR, presuming the transmission power given by the MN’s hardware configuration and a desired target payload rate. For simulating the channel characteristics faced by the MN, the model comprises parameters like the path loss in dependence of the signal-center frequency, the transmission distance and receiver and transmitter heights. Environmental influences like attenuation by obstacles, multi-path propagation or unpredictable rotation of the MN’s rod antenna are incorporated by means of a random variable, stating the superimposed attenuation effects. Based on these assumptions we were capable of overcoming path losses of over 150dB for a distance of 5 km and more, under reasonable rates of packet loss^17^, thus accomplishing an ultra-robust implementation supporting payload data rates of a few bits per second.

### Sensor node energy consumption and runtime

A crucial aspect for biologging is knowledge on the runtime of the sensor nodes. Static program-code analysis methods of the mobile node are able to determine upper bounds on the nodes’ runtime^48^. However, in the context of the BATS tracking system, precise estimates for the average uptimes of the system proved to be more beneficial for the empirical studies than upper bounds for the lifetime. Consequently, we focused on an energy model to determine the average runtime of the mobile nodes, which is strongly dependent on the tasks executed by the software. Our models are based on measurements of each executed task in combination with empirically determined activity parameter of each task. That way, we ensure highest accuracy for our model. In our setting, seven different tasks are implemented: (i) Standby, (ii) sending MN-beacons, (iii) receiving MN-beacons, (iv) observing ground node availability, (v) transmitting data to a ground node, (vi) sending localization packets, and (vii) sending presence signals (see Table S Glossary for definition of terms).

We determined the runtime by using the specific energy demand for a task and by translating it to an average current draw. With the average current draw and a given battery capacity, the runtime can be computed as follows:

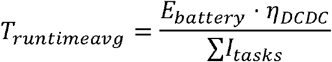

ŋDCDC represents the efficiency of the DCDC-converter, which is permanently active and consumes energy. Determining ŋDCDC is impractical, because it highly depends on the actual current drain of the application for the entire runtime. For this reason, we assume a fixed efficiency of 0.95, which translates to only 95% of the battery capacity being available for software tasks. This way, losses caused by the DCDC and parasitic discharges of the battery are modeled in a coarse-grained manner.

The idle current during standby is given in a current draw, which does not require any further calculations. The other tasks (e.g., observing GN availability, sending localization packets) are executed in predefined cycle times (duty cycle). Based on the measured energy demand and the duty cycle, we calculated an average current draw for each task. The energy demand for each task was measured in the lab with an Agilent DC power analyzer precise source meter. In the case of a localization packet, which is sent every 128ms, the average current draw can be expressed as follows:

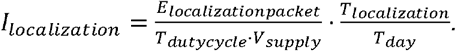

The average time spent inside the localization grid *(T*_*localization*_) per day is strongly dependent on the species-specific animal behavior and the experimental design. For the calculations presented here, we set the daily localization period to 2 or 4 hours, respectively.

Observing a ground node in receiving range is carried out at a fixed duty cycle of 2 seconds. Here, the task is independent of the behavior of the tracked animal and the energy demand is calculated as follows:

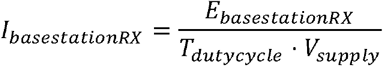

The transmission rate of MN-beacons and presence signals is adaptive based on the contact to a ground node (i.e., a mobile node near a GN at a roost will decrease the duty cycle in comparison to a MN on a foraging animal which is not in reception range of a GN). In turn, the energy demand for sending beacons and presence signals highly depends on the behavior of the tracked study species and the individual animal (e.g., time spent near GNs at roosting sites). We therefore used empirical data obtained in the Berlin field test to inform our energy model with realistic averaged parameter values for duty cycles of each task and the amount of transmitted data. We quantified the average time-tagged bats spent in reception range of a GN (inactive mode, decreased duty cycle) versus the time bats spent outside the reception range of any GN (active mode, increased duty cycle). The common noctule bats in the Berlin field test spent on average 47% of the observation time in the inactive mode and the energy demand for transmitting beacons and presence signals calculates as follows:

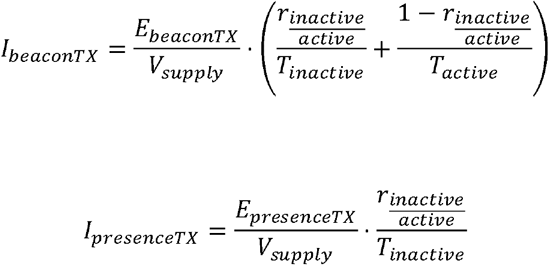

Receiving a MN-beacon depends on the duty cycle at which beacons are transmitted and on the number of mobile nodes in receiving range. During the Berlin field test, we calculated the average number of 2.05 maximum parallel meetings.

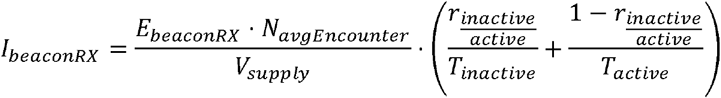

For data download to a GN, we assumed static energy consumption (the energy demand for sending a data packet highly depends on the size of the packet to be transmitted^23^. The number of data packets to be transmitted is again dependent on the behavior of the tracked animal (depending on how many meetings an individual accumulates). Mobile nodes on common noctule bats sent on average 23.7 packages per hour to a ground node. Thus the current draw can be denoted as:

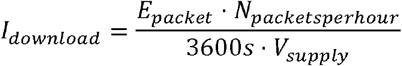

Based on these calculations, we matched the estimated average runtimes to the observed runtimes during the Berlin field test (based on the last beacon or packet received from each individual tagged bat).

### Adaptive operation, scalability, and reconfiguration

The adaptive operation contributes to the energy efficiency of BATS. We define location-specific communication schemes on the mobile nodes which are initiated by ground nodes. At a noctule bat day roost, for example, GNs activated the inactive beacon interval where MN-beacons for meeting generation were only sent every 10 seconds. When tagged bats leave their roost and move beyond the reception range of the GN, the MN switches to the active interval, sending a MN-beacon every two seconds, which increases the probability to detect also very short meetings in comparison to the inactive rate. Similarly, localization packets, which strongly increase the energy demand, should only be sent when the tagged animal moves within the tracking grid and are therefore triggered by a GN within the grid.

BATS tracks multiple individuals simultaneously. Our current design allows for the observation of up to a theoretical maximum of 60 individuals. Field deployments containing 11 to 50 tagged bats empirically validated this targeted scalability. The scale of the localization grid can be adapted to the ranges required for experimental setups. While we tracked mouse-eared bats on 1.5ha, smaller areas might be sufficient to track, e.g., rodents (bank voles, which showed a density of 40 - 162 individuals per hectare, have been tracked on less than 0.5ha using automated VHF telemetry^49^). Tracking grids larger than 1.5ha are certainly possible from a technological point of view. Yet, one has to keep in mind that effort for maintenance (e.g., replacing power sources for localization nodes) scales with the size of the tracking grid.

Since settings for communication schemes are sometimes difficult to pick a priori, we built an option for reconfiguration of so-called ‘soft-settings’ (active/inactive rate of MN-beacons, RSSI-thresholds for data download, localization interval, or timeout duration after which a running meeting is terminated, etc.). Four values for every soft-setting (e.g., active rate 2s, 4s, 10s, 30s) can be defined a priori and during operation ground nodes can be used to trigger a switch between these pre-defined values at the mobile node.

### Field deployments

BATS functions as a modular system and hardware setup and software configurations are easily tailored to a specific use case. We evaluated BATS during three major field studies by applying mobile nodes to vampire bats, noctule bats, and mouse-eared bats with body weights of 27 - 48g, 18 - 35g, and 22 - 28g respectively. While at least two of the three major functionalities (proximity sensing, high-resolution tracking, long-range telemetry) have been used in all three field studies, we focus on one specific functionality per deployment.

### Proximity sensing in vampire bats

We tagged 50 common vampire bats (*Desmodus rotundus;* 44 adult females, 6 subadults) from a colony roosting in a cave tree near Tolé, Panama, to document social networks with high resolution. Field work was conducted during September and October 2017. Mobile nodes were powered by a 22mAh LiPo battery and housed in a 3D-printed plastic case, resulting in a total weight of 1.8g. One download-dedicated ground node was positioned inside the roost and five ground nodes were placed on surrounding cattle pastures to detect the presence of foraging or commuting bats.

### Long-range telemetry in noctule bats

We captured and tagged 34 common noctule bats (*Nyctalus noctula*; 19 adult females and 15 juveniles) from two bat boxes in a nursing colony an urban forest in the city of Berlin, Germany (‘Königsheide Forst’)^50^. Mobile nodes were powered by either a 12mAh or a 22mAh battery and were housed either in a 3D-printed plastic case or in a fingertip of a nitrile lab glove that was sealed with glue. Total weight varied between 1.1 - 1.9g depending on housing and battery. We positioned 5 ground nodes underneath known roosts to document the presence of individual bats and to remotely download data. In addition, we set up two long-range receivers to evaluate model-based predicted data retrieval over distances of up to 4-5km^17^. This opportunity is particularly valuable to retrieve data of tagged individuals that moved to an unknown roost.

### High-resolution tracking of mouse-eared bats

We captured 11 mouse-eared bats (*Myotis myotis*) using mist-nets set up at ground level in a mature deciduous forest near Forchheim, Germany. When hunting for ground beetles, mouse-eared bats are faithful to their foraging sites for consecutive days. We therefore mist-netted bats at an attractive foraging site rather than catching them from a roost in order to track repeated bouts by returning individuals over the course of several days. Mobile nodes were powered by 22mAh batteries and housed in fingertips of nitrile lab gloves (total weight 1.4g). At the capture site we installed a tracking grid consisting of 17 localization nodes covering roughly an area of 1.5ha (see Figure 3). Distance between tracking stations varied between ca. 25 - 40m. The irregular configuration was due to the presence of thick trees. We aimed at positioning tracking stations at least 3 - 5m away from trees to reduce shielding of the signal. We set up a polygon-shaped reference path for estimating localization errors and determined the true position of the corners using a Leica Robotic Total Station TS16 (positioning error < 5cm). Corners were connected using strings, and we walked either a sensor node (once) or a GPS tracker (four times) (Ornitela OrniTrack-15; a 15g solar powered GPS-GSM/GPRS tracker; maximum logging rate 1 fix per second at a lifetime of ca. 4h without solar harvesting) along the calibration path and calculated the average localization error based on the obtained tracks.

### Ethics

#### Work on vampire bats

All experiments were approved by the Smithsonian Tropical Research Institute Animal Care and Use Committee (#2015-0915-2018-A9 and #2017-0102-2020) and by the Panamanian Ministry of the Environment (#SE/A-76-16 and #SE/AH-2-17).

#### Work on noctule bats

All necessary permits were obtained from SenStadtUm (I E 222/OA-AS/G_1203) and LaGeSo (I C 113-G0008/16).

#### Work on mouse-eared bats

All experiments were approved by the government of Upper Franconia (55.1-8642.01-15/13) and by the government of Lower Franconia (55.2-DMS-2532-2-181).

## 5. Acknowledgements

This study was funded by grants of the Deutsche Forschungsgemeinschaft (FM, AK, RK, KMW, WSP, JT, JR, FD) within the research unit FOR-1508, a Smithsonian Scholarly Studies Awards grant (RAP, GGC, SPR, FM), and a National Geographic Society Research Grant WW-057R-17 (GGC). We thank M Mutschlechner, M Nabeel, J Blobel, and M Hierold for their contributions within the scope of the research unit. We highly appreciate logistical support during the Forchheim field study by Naturstrom, MTS Meixner Transporte + Service, C. Kreul GmbH & Co. KG, Spedition Pohl GmbH & Co. KG, and Siedlergemeinschaft Lichteneiche. We are grateful to J Berrío-Martínez, D Josic, M Nowak, G Cohen, L Günther, H Wieser, C Rüstau, B Peiffer, M Hammer, F Oehme and his BUND group of volunteers for their support during field work. We express our particular thanks to J and C Mohr, who have substantially contributed to the success of our field work. We thank E. Siebert for making the line work of *D. rotundus*.

## Author contributions

SPR and FM conceived the ideas and designed the biologging studies; ND, BC, MH, TN, PW, and MS developed and tested the tracking technology; SPR, GGC, ND, BC, MH, TN, and MS collected the data; MH, TN, MS, SH, BC, PW and SPR analyzed the data; SPR led the writing of the manuscript. SPR, GGC, RAP, AK, RW, JT, JR, KMW, WSP, RK, FD, and FM contributed to acquiring funds. All authors contributed critically to the drafts, gave final approval for publication and agree to be held accountable for the work performed therein.

---

**Supplementary Figure 1:**
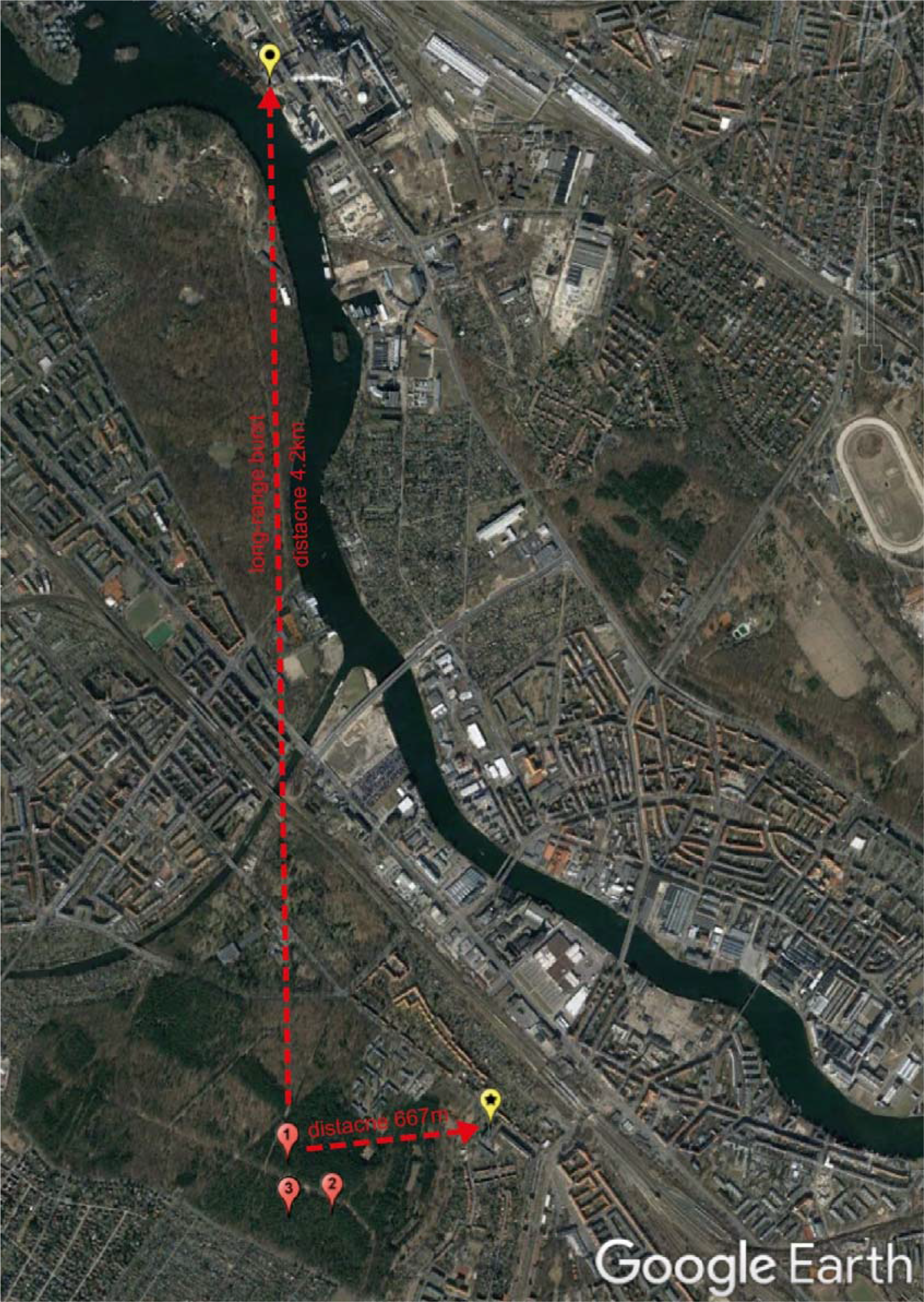
Experimental setup for long-range telemetry in the city of Berlin, Germany. Yellow symbols mark the positions of two long-range telemetry receivers (asterix = retirement home, circle = cogeneration plant). Red symbols show the locations of bat day roosts 1 to 3 (tree holes). Map data ©2019 GeoBasis-DE/BKG (©2009), Google.

**Supplementary Figure 2:**
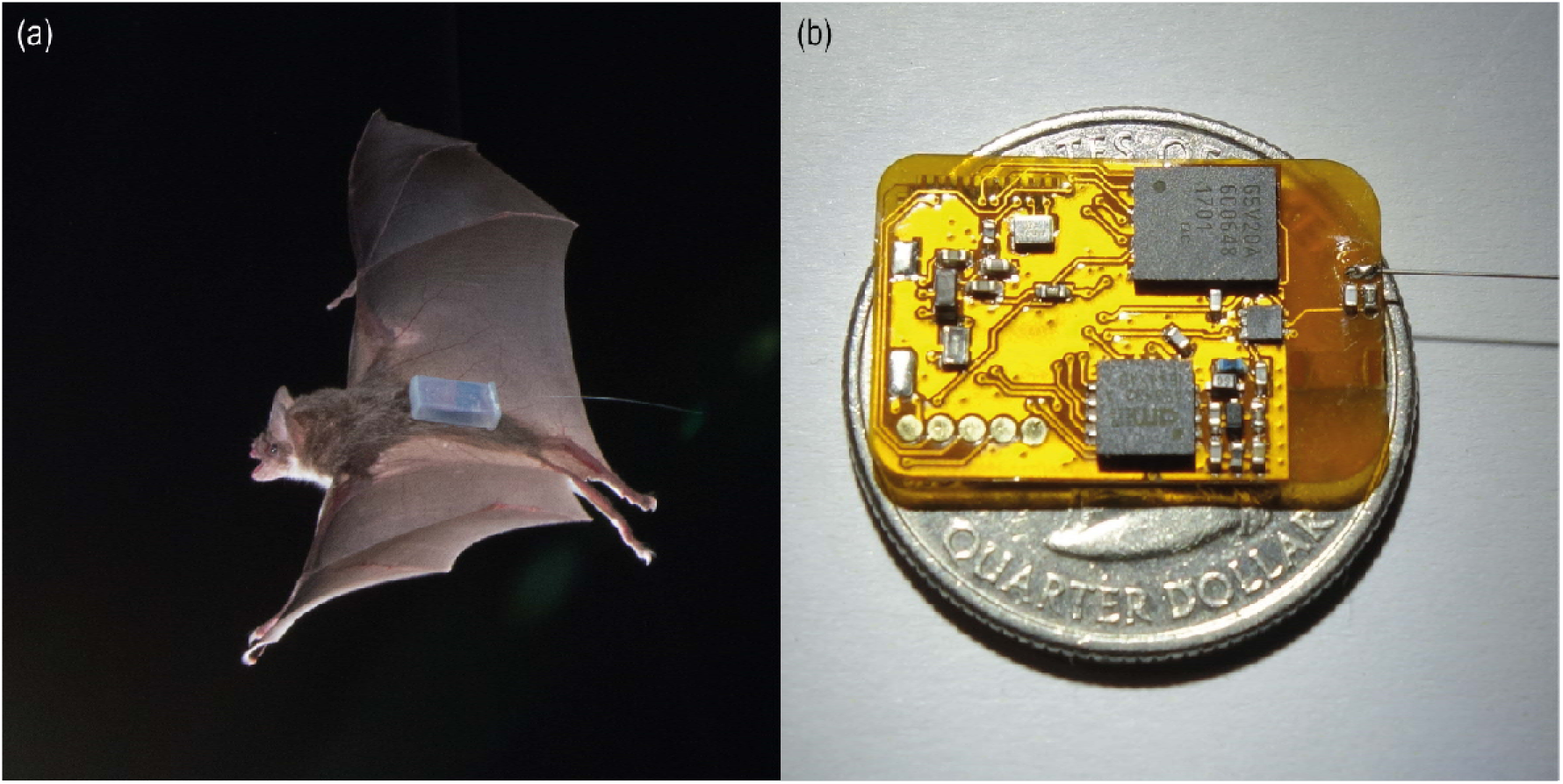
animal-borne mobile node for proximity sensing. (a) Common vampire bat (*Desmodus rotundus*) carrying a mobile node housed in a plastic case; (b) bare mobile node on a quarter US dollar coin for comparison of size.

**Supplementary Table 1:** Glossary. Description of hardware components, communication and data types and software tasks considered for the energy model.

| Keyword | Description |
| --- | --- |
| **Hardware** |  |
| Mobile node (MN) | Animal-borne sensor node. |
| Ground node (GN) | Multifunctional ground-based sensor nodes. Depending on the GN-configuration, it receives data from MNs, triggers transmission of localization packets at the MN, or triggers the transmission of presence signals at the MN. |
| Localization node | Sensor node containing a dual-band antenna for high-resolution positioning of MNs. Arranged in a tracking grid. |
| Long-range receiver | Long-range telemetry-dedicated sensor node for receiving long-range bursts. |

| **Communication and data types** |  |
| --- | --- |
| Ground node beacon<br>(GN-beacon) | A packet sent by a ground node that triggers action on the MN (e.g., data download). |
| Mobile node beacon<br>(MN-beacon) | A packet sent by a mobile node. It contains a wakeup sequence which wakes other MNs from standby. |
| Long-range burst | A packet embedded in the wakeup sequence of the MN-beacon for data transmission to a long-range receiver. |
| Localization packet | A packet sent by a MN to localization nodes for high-resolution tracking. Transmission is initiated by a GN-beacon. |
| Presence Signal | A packet sent by a MN to indicate the presence of a bat in range of a GN. Transmission is initiated by a GN-beacon. |
| **Software task on the mobile node** |  |
| (i) standby | An idle state where only mandatory devices are active (e.g., wakeup receiver). The idle state is terminated by the reception of a wakeup. |
| (ii) sending beacons | Sending a MN-beacon to wake nearby MNs from standby. Nearby MNs immediately send their IDs in return. |
| (iii) receiving beacons | If the wakeup receiver receives a wakeup sequence, the conventional receiver is activated to receive the IDs of nearby MNs (encoded in the MN-beacons). |
| (iv) observing ground node availability | Every 2 s the receiver is activated to check whether GN-beacons are received. |
| (v) transmitting data to a ground node | If beacons of a download-dedicated GN are received, the transmitter is activated and data download is initiated. |
| (vi) sending localization packets | If beacons of a tracking-dedicated GN are received, localization packets are sent (duty cycle 8/s; 868MHz & 2.4GHz) |
| (vii) sending presence signals | If beacons of a GN are received, a presence signals is send |

## References

1. Kays, R., Crofoot, M.C., Jetz, W. & Wikelski, M. Terrestrial animal tracking as an eye on life and planet. Science 348, aaa2478 (2015).

2. Krause, J. et al. Reality mining of animal social systems. Trends Ecol. Evol. 28, 541–551 (2013).

3. Wilmers, C.C. et al. The golden age of bio-logging: how animal-borne sensors are advancing the frontiers of ecology. Ecology 96, 1741–1753 (2015).

4. Hughey, L.F., Hein, A.M., Strandburg-Peshkin, A. & Jensen, F.H. Challenges and solutions for studying collective animal behaviour in the wild. Philosophical Transactions of the Royal Society B: Biological Sciences 373, 20170005 (2018).

5. McKinnon, E.A. & Love, O.P. Ten years tracking the migrations of small landbirds: Lessons learned in the golden age of bio-logging. The Auk 135, 834–856 (2018).

6. Fraser, K.C. et al. Tracking the conservation promise of movement ecology. Frontiers in Ecology and Evolution 6, 150 (2018).

7. Curry, A. The internet of animals that could help to save vanishing wildlife. Nature 562, 322 (2018).

8. Oppermann, F.J., Boano, C.A. & Römer, K. in The Art of Wireless Sensor Networks 11–50 (Springer, 2014).

9. Juang, P. et al. Energy-efficient computing for wildlife tracking: design tradeoffs and early experiences with ZebraNet. SIGPLAN Not. 37, 96–107 (2002).

10. Toledo, S., Kishon, O., Orchan, Y., Shohat, A. & Nathan, R. in Software Science, Technology and Engineering (SWSTE), 2016 IEEE International Conference on 51-60 (IEEE, 2016).

11. Rutz, C. et al. Automated mapping of social networks in wild birds. Curr. Biol. 22, R669–R671 (2012).

12. Ayele, E.D., Meratnia, N. & Havinga, P.J. in New Technologies, Mobility and Security (NTMS), 2018 9th IFIP International Conference on 1-5 (IEEE, 2018).

13. Ripperger, S. et al. Automated proximity sensing in small vertebrates: design of miniaturized sensor nodes and first field tests in bats. Ecology and Evolution 6, 2179–2189 (2016).

14. Wilkinson, G.S. et al. Kinship, association, and social complexity in bats. Behav. Ecol. Sociobiol. 73, 7 (2019).

15. Sekara, V., Stopczynski, A. & Lehmann, S. Fundamental structures of dynamic social networks. Proceedings of the national academy of sciences 113, 9977–9982 (2016).

16. Kilian, G. et al. Increasing Transmission Reliability for Telemetry Systems Using Telegram Splitting. IEEE Transactions on Communications 63, 949–961 (2015).

17. Schadhauser, M., Robert, J. & Heuberger, A. in Smart SysTech 2017; European Conference on Smart Objects, Systems and Technologies 1-8 (VDE, 2017)

18. Hallworth, M.T. & Marra, P.P. Miniaturized GPS tags identify non-breeding territories of a small breeding migratory songbird. Scientific reports 5, 11069 (2015).

19. Cvikel, N. et al. Bats Aggregate to Improve Prey Search but Might Be Impaired when Their Density Becomes Too High. Curr. Biol. 25, 206–211 (2015).

20. Conenna, I., López-Baucells, A., Rocha, R., Ripperger, S. & Cabeza, M. Movement seasonality in a desert-dwelling bat revealed by miniature GPS loggers. Movement ecology 7, 1–10 (2019).

21. Tomkiewicz, S.M., Fuller, M.R., Kie, J.G. & Bates, K.K. Global positioning system and associated technologies in animal behaviour and ecological research. Philosophical Transactions of the Royal Society B: Biological Sciences 365, 2163–2176 (2010).

22. Dressler, F. et al. Monitoring Bats in the Wild: On Using Erasure Codes for Energy-Efficient Wireless Sensor Networks. ACM Transactions on Sensor Networks (TOSN) 12, 1–29 (2016).

23. Cassens, B. et al. in Proceedings of the 2019 International Conference on Embedded Wireless Systems and Networks 59–70 (Junction Publishing, 2019).

24. St Clair, J. et al. Experimental resource pulses influence social-network dynamics and the potential for information flow in tool-using crows. Nature Communications 6, 7197 (2015).

25. O’Mara, M.T., Wikelski, M. & Dechmann, D.K. 50 years of bat tracking: device attachment and future directions. Methods in Ecology and Evolution 5, 311–319 (2014).

26. Duda, N. et al. in 2019 IEEE Topical Conference on Wireless Sensors and Sensor Networks (WiSNet) 1–4 (IEEE, 2019).

27. Kays, R. et al. Tracking Animal Location and Activity with an Automated Radio Telemetry System in a Tropical Rainforest. The Computer Journal (2011).

28. Nowak, T., Hartmann, M., Tröger, H.-M., Patino-Studencki, L. & Thielecke, J. in 2017 IEEE International Conference on Communications Workshops (ICC Workshops) 1024–1029 (IEEE, 2017).

29. Kerth, G., Perony, N. & Schweitzer, F. Bats are able to maintain long-term social relationships despite the high fission–fusion dynamics of their groups. Proceedings of the Royal Society B: Biological Sciences 278, 2761–2767 (2011).

30. Aplin, L.M. et al. Experimentally induced innovations lead to persistent culture via conformity in wild birds. Nature 518, 538–541 (2015).

31. Lopes, P.C., Block, P. & König, B. Infection-induced behavioural changes reduce connectivity and the potential for disease spread in wild mice contact networks. Scientific reports 6, 31790 (2016).

32. Levin, I.I. et al. Stress response, gut microbial diversity and sexual signals correlate with social interactions. Biol. Lett. 12, 20160352 (2016).

33. Saltzer, J.H., Reed, D.P. & Clark, D.D. End-to-end arguments in system design. ACM Transactions on Computer Systems 100, 277–288 (1984).

34. Schmidt, A. et al. Cross-Layer Pacing for Predictably Low Latency. Proc. 6th Intl. Worksh. on Ultra-Low Latency in Wireless Networks (Infocom ULLWN). IEEE (2019).

35. Blobel, J. & Dressler, F. in 2017 IEEE Conference on Computer Communications Workshops (INFOCOM WKSHPS) 984–985 (IEEE, 2017).

36. Nabeel, M., Amjad, M.S. & Dressler, F. in 2018 IEEE Global Communications Conference (GLOBECOM) 1–6 (IEEE, 2018).

37. Teitelbaum, C.S. & Mueller, T. Beyond Migration: Causes and Consequences of Nomadic Animal Movements. Trends Ecol. Evol. (2019).

38. Stroeymeyt, N. et al. Social network plasticity decreases disease transmission in a eusocial insect. Science 362, 941–945 (2018).

39. Consortium, T.Q. Networking Our Way to Better Ecosystem Service Provision. Trends Ecol. Evol. 31, 105–115 (2016).

40. Nowak, T., Hartmann, M., Zech, T. & Thielecke, J. in 2016 IEEE-APS Topical Conference on Antennas and Propagation in Wireless Communications (APWC) 110–113 (IEEE, 2016).

41. Nowak, T., Hartmann, M. & Thielecke, J. Unified performance measures in network localization. EURASIP Journal on Advances in Signal Processing 2018, 48 (2018).

42. Hartmann, M., Nowak, T., Pfandenhauer, O., Thielecke, J. & Heuberger, A. in 2016 12th Annual Conference on Wireless On-demand Network Systems and Services (WONS) 1–8 (IEEE, 2016).

43. Nowak, T., Hartmann, M. & Thielecke, J. in 2018 11th German Microwave Conference (GeMiC) 115–118 (IEEE, 2018).

44. Nowak, T., Hartmann, M., Patino-Studencki, L. & Thielecke, J. in 2016 13th Workshop on Positioning, Navigation and Communications (WPNC) 1–6 (IEEE, 2016).

45. Duda, N. et al. BATS: Adaptive Ultra Low Power Sensor Network for Animal Tracking. Sensors 18, 3343 (2018).

46. Hijmans, R.J., Williams, E., Vennes, C. & Hijmans, M.R.J. in Spherical trigonometryv 1.5–7 (2019).

47. Schadhauser, M., Robert, J. & Heuberger, A. in Smart SysTech 2016; European Conference on Smart Objects, Systems and Technologies 1–9 (VDE, 2016).

48. Wägemann, P., Dietrich, C., Distler, T., Ulbrich, P. & Schröder-Preikschat, W. Whole-system worst-case energy-consumption analysis for energy-constrained real-time systems. Leibniz International Proceedings in Informatics, LIPIcs 106 (2018) 106, 24 (2018).

49. Schirmer, A., Herde, A., Eccard, J.A. & Dammhahn, M. Individuals in space: personality-dependent space use, movement and microhabitat use facilitate individual spatial niche specialization. Oecologia 189, 647–660 (2019).

50. Ripperger, S. et al. Proximity sensors on common noctule bats reveal evidence that mothers guide juveniles to roosts but not food. Biol. Lett. 15, 20180884 (2019).

